# Multiple Wnt signaling pathways direct epithelial tubule interconnection in the regenerating zebrafish kidney

**DOI:** 10.1101/2025.03.26.645545

**Authors:** Caramai N. Kamei, William G. B. Sampson, Carolin Albertz, Oliver Aries, Amber Wolf, Rohan M. Upadhyay, Samuel M. Hughes, Heiko Schenk, Frederic Bonnet, Bruce W. Draper, Kyle W. McCracken, Denise K. Marciano, Leif Oxburgh, Iain A. Drummond

## Abstract

Epithelial tubule fusion is fundamental for kidney morphogenesis. Differentiating nephron tubules interconnect with collecting system epithelia to generate a lumenal pathway for fluid excretion. In the adult zebrafish kidney, nephrogenesis occurs as a regenerative response to injury and provides a model to explore cell signaling pathways required for tubule interconnection. We show that canonical Wnt signaling at the junction between two tubules induces a mesenchymal, invasive cell phenotype and is required, along with Src kinase and rac1, to generate basal cell protrusions. The Wnt ligands *wnt9b* and *wnt4* are both required for new nephron formation after injury. Mutation in *wnt4* or treatment with the canonical Wnt inhibitor IWR1 blocks formation of basal protrusions in forming nephrons. Mutation in the Wnt receptor *frizzled9b* reveals a fusion-associated non-canonical Wnt pathway that acts to 1) restrict canonical Wnt gene expression, 2) drive Rho kinase-dependent apical constriction of epithelial cells, and 3) position basal protrusions and generate orthogonal tubule lumenal connections. As a result, *frizzled9b* mutant nephrons fail to fully interconnect with target distal tubules. Our results indicate that canonical and non-canonical Wnt signaling interact in the same cells to orient and drive tubule interconnection in the regenerating zebrafish kidney.

## Introduction

Embryonic development and epithelial organogenesis require fusion of cell sheets or tubules to create polarized transport and secretory structures that are critical for tissue function ^1^. Epithelial fusion is a complex process involving regionally specialized cells that partially uncouple from cell neighbors to undergo an incomplete mesenchymal transformation with the formation of invasive or exploratory actin-based cell protrusions ^2^. These transient changes in cell adhesion and polarity allow for cell rearrangements and integration with new cell neighbors followed by resolution of cell junctions, stabilization of cell adhesion, and restoration of a uniform epithelial cell polarity ^2^. Examples of epithelial sheet fusion include the morphogenetic process of neural tube closure ^3^, optic cup fusion in the developing eye ^4,5^, and palate fusion ^6^. These fusion events require cell movement, lamellipodia extension, initiation of new cell adhesions, and finally stabilization of uniform cell polarity. The fusion of two epithelial tubules has the added requirement of invasion into tubule basement membranes, as basal to basal cell apposition must be overcome to allow tubules to join lumens ^6,7^. Currently, the cell signaling processes that mediate vertebrate tubule fusion are not well understood.

Epithelial tubule fusion is observed during development of the Drosophila tracheal system ^8^, the vertebrate pancreas ^9^, and in the avian lung ^1^. In Drosophila, FGF signaling plays a prominent role in guiding fusion of tracheal tip cells and supporting lumenal interconnection between single cell thick tubules ^10^. In the chick lung, an elongating branched multicellular epithelium forms a continuous network of airways by a mass fusion event involving changes in epithelial shape at apposing tubule tips, formation of bridging cytoskeletal protrusions, removal of intervening basement membrane, and cell apoptosis at tubule junctions, making a path for lumen fusion ^11^. Signals driving avian epithelial budding and fusion are currently not known.

In the embryonic mammalian kidney, epithelial tubule fusion is required for the integration of newly forming epithelial nephron tubules into an arborized network of filtration units ^7,12^. This process is characterized by a thinning of tubule basement membranes, invasion of new nephron cell basal protrusions, and formation of a connecting lumen between the distal end of S-shaped body nephron intermediates and the branched tips of the ureteric epithelium that will form the collecting system ^7,13,14^. This fusion event requires distal patterning of the invading S-shaped body, implying that invasive, interconnecting cell behavior is a differentiated phenotype of the distal tubule ^7^. Identifying signaling mechanisms that drive nephron fusion will be essential as a critical step in stem cell-derived kidney organoid engraftment to host kidneys and to develop kidney organoids for augmenting renal function ^15–18^.

In the adult zebrafish kidney, nephron addition and tubule fusion occur throughout life as the kidney grows ^19–21^. Nephron addition also occurs in response to acute kidney injury where adult renal progenitor cells migrate to form rosette-like aggregates on distal tubules prior to new nephron outgrowth, patterning, and interconnection ^22,23^. We have shown that cell aggregation and rosette formation require FGF signaling ^22^ while tubule outgrowth requires canonical Wnt signaling ^23^. Here we demonstrate that the final step of new nephron integration requires both canonical and non-canonical Wnt signals acting in a single layer of invading tubule cells. We also identify downstream morphogenetic processes involved in the generation of Src and Rac1-dependent basal protrusions and show that non-canonical *frizzled9b* signaling is required to orient invasive processes and coordinate apical and basal constriction of invading tubule cells. Our results illuminate signaling events in normal nephron tubule interconnection and also suggest how cell signaling could be engineered to direct engraftment of kidney organoid-derived nephrons, currently a bottleneck in kidney regenerative medicine.

## Results

### Regenerating new kidney nephrons form lumenal connections by invasion of existing distal tubules

We took advantage of the synchronous addition of up to 100 new nephrons at 7-8 days post-acute kidney injury ^24^ to dissect cell signaling and morphogenetic processes guiding tubule fusion. Zebrafish kidney progenitor cells marked by *Tg(lhx1a:egfp)* expression ^21^ and kidney tubule lumens visualized by the ring of apical actin at the lumenal surface were imaged before (figure 1A; supplemental movie 1) and after tubule fusion (figure 1B; supplemental movie 2), 7-8 days following gentamicin-induced acute kidney injury. New nephron epithelial cells are apically constricted prior to fusion (figure 1A,C,E) and transition to a basally constricted morphology during the process of tubule fusion (figure 1D,F). Preceding interconnection of new and existing nephron tubules, the basal surfaces of newly formed *lhx1a:eGFP+* nephrons adjacent to a distal tubule segment exhibit multiple basal protrusions that invade the target distal tubule, preceding interconnection of new and existing tubule lumens (figure 1G; supplemental movie 3). These basal protrusions are composed of a central core of filamentous actin and form at the periphery of the rosette as well as directly under the forming lumen of new nephrons (figure 1A inset, C).

**Figure 1.**
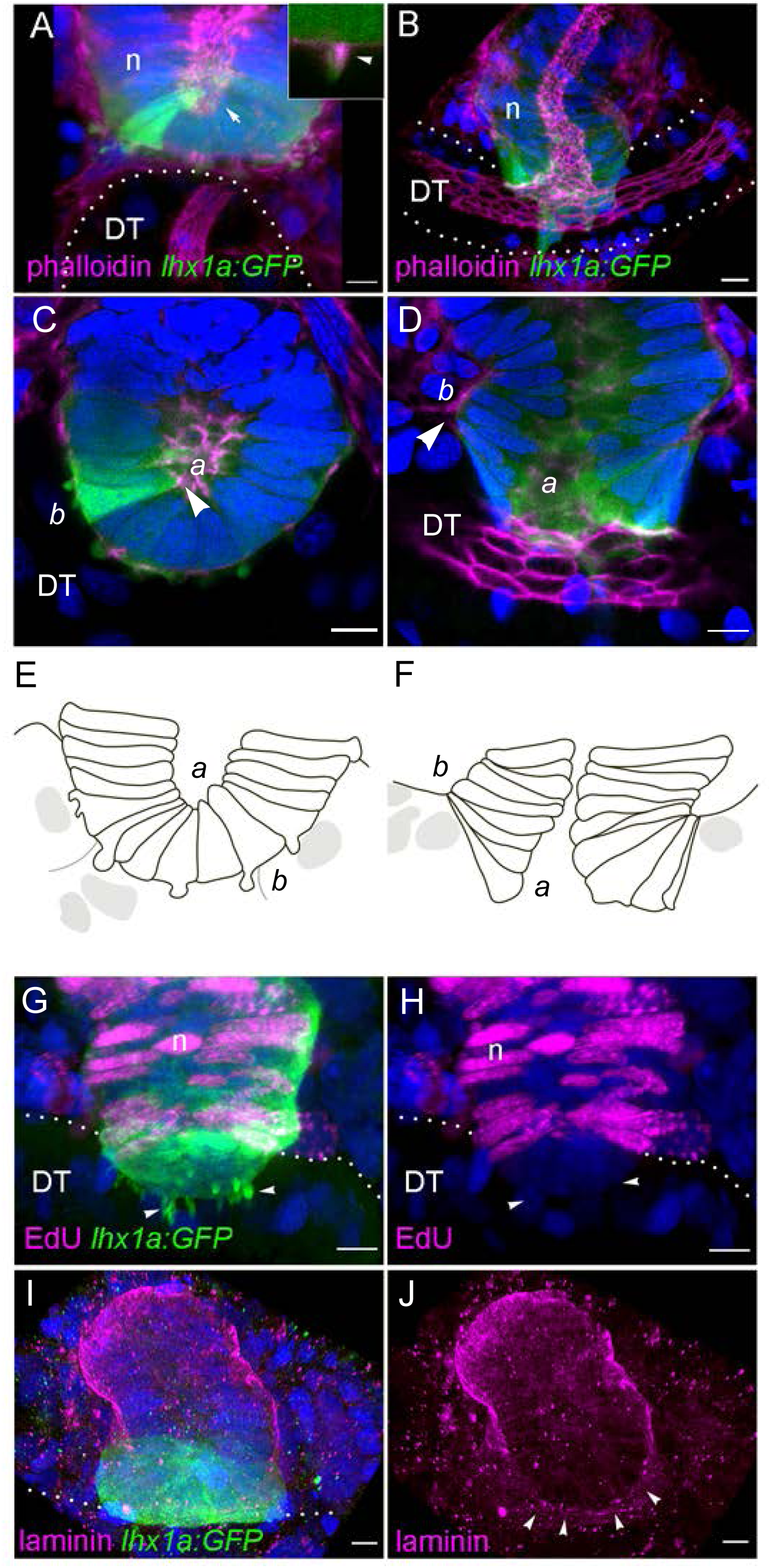
Stages of epithelial tubule interconnection. **(A)** Prior to tubule fusion phalloidin staining (magenta) of apical cell actin highlights a new tubule epithelial lumen (arrow) in the process of nephron fusion to the distal tubule (DT; dotted outline). *Tg(lhx1a:egfp)* (green) marks newly forming new nephrons (n) adjacent to target distal tubules (DT). Junctional nephron cells are constricted apically. **(A, inset)** Closeup of basal protrusion with phalloidin positive actin core. **(B)** After tubule fusion, phalloidin staining (magenta) outlines the apical cell actin and lumen continuity between old and new nephron tubules. (**C**; *a*, apical; *b*, basal) but become basally constricted in the process of lumen fusion (**D**). Diagram of cell boundaries drawn from C, D **(E, F)**. Basal protrusions (G, arrowheads) invade the distal tubule at the point of tubule fusion; these cells are non-proliferating (H; arrowheads). New nephrons lack laminin-positive basement membranes at their basal surface point of fusion marked by *Tg(lhx1a:egfp)* expression (H,I; arrowheads). Blue fluorescence is Hoechst-stained nuclei. Scale bars = 5 µm except for B where scale bar = 10 µm.

New nephron cells closest to the target distal tubule do not proliferate as revealed by the lack of EdU uptake in invading cells (figure 1H). Proliferating, EdU-positive cells in the new nephron extend the nascent tubule ^23^ and develop a laminin-containing basement membrane typical of epithelial cells while invasive cells closest to the point of future interconnection are laminin-negative (figure 1I,J). To confirm the observed invasive phenotype with a molecular signature, we compared a prior microarray transcriptome of *Tg(lhx1a:egfp)+* cells with the gene set represented in the human cancer metastasis database ^21,25^. Zebrafish orthologs of genes associated with metastatic cell invasive behavior were expressed at points of nephron connection including invadopodia-associated metalloproteases *mmp14a, mmp14b,* the SH3 domain protein *tks5*, the Wnt ligand *wnt4*, canonical Wnt targets *lef1* and *cdh11* implicated in invasive cell behavior, and transcriptional regulators of invasiveness, *id1* and *cjun* ^25,26^ (Supplemental figure 1). Taken together, the data indicate that a narrow band of new nephron cells at the point of tubule fusion maintain a non-proliferating, invasive mesenchymal character, while new nephron cells further from the point of interconnection develop an epithelial basement membrane and proliferate, extending the nascent epithelial tubule.

### Canonical Wnt signaling stimulates tubule invasion

Wnt signaling is required for zebrafish new nephron outgrowth and Wnt ligands *wnt9a* and *wnt9b* are induced after acute kidney injury specifically in distal tubules where new nephrons form ^23^. To better characterize expression domains of known canonical Wnt signaling ligands and targets, we performed confocal 3D imaging of *wnt4* and *lef1* expression by fluorescent in situ hybridization. Both *wnt4* and *lef1* were expressed in a restricted pattern in cells at the point of tubule fusion (figure 2A, B), consistent with high canonical Wnt signaling in this domain ^27,28^. The zebrafish canonical Wnt reporter *Tg(tcflef:dGFP)* also showed highest expression at the point of contact with the distal tubule (figure 2C; ^29^).

**Figure 2.**
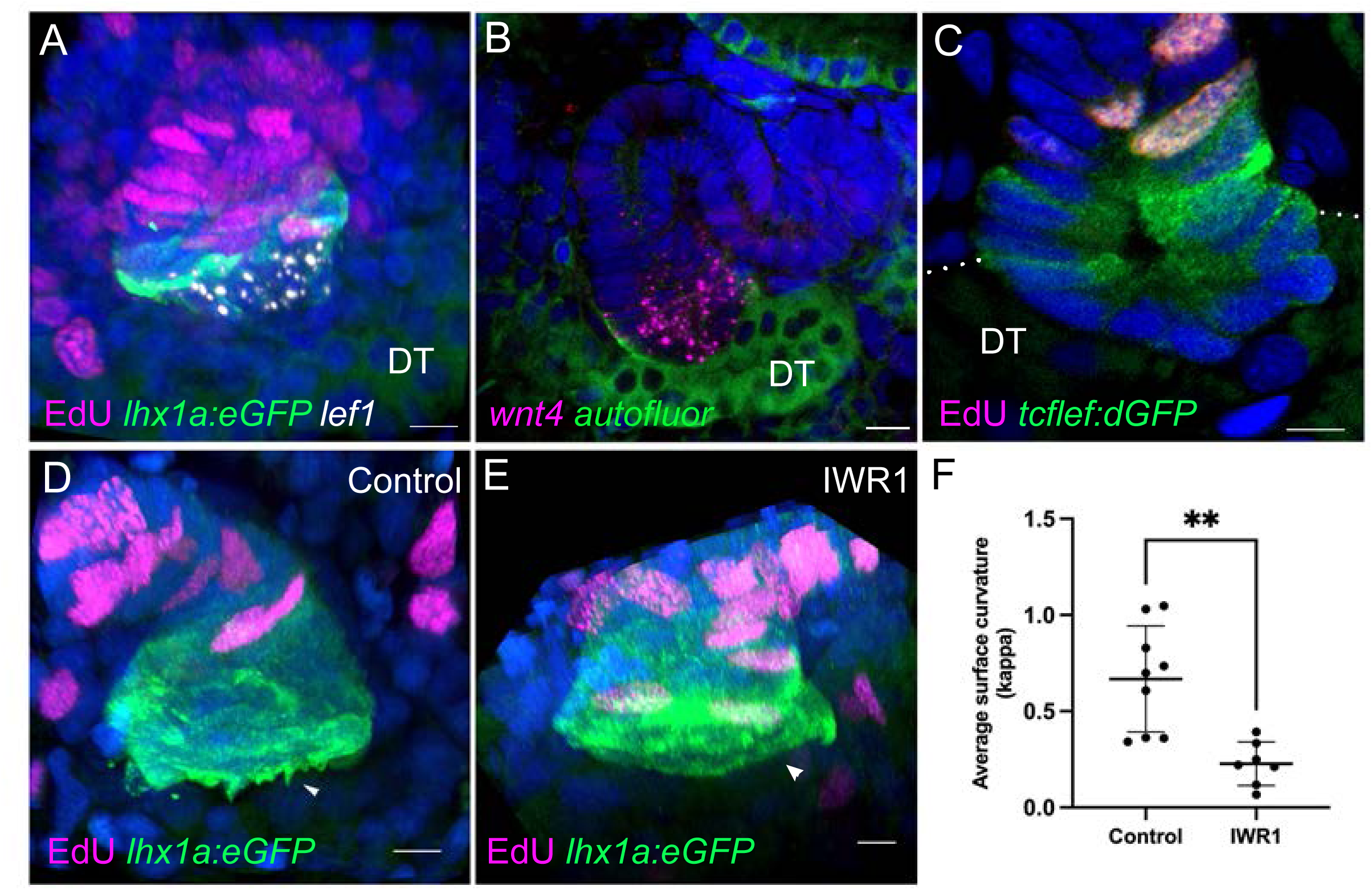
Canonical Wnt signaling and tubule invasion. Canonical Wnt target genes *lef1* **(A)** and *wnt4* **(B)** are expressed in new nephron tubules (marked by EdU incorporation, at the tubule fusion junction marked by *Tg(lhx1a:egfp)* expression (**A,B)**. The *Tg(tcflef:dGFP)* Wnt reporter transgene is expressed at the point of tubule fusion (C). The canonical Wnt inhibitor IWR1 blocks formation of invasive basal protrusions (arrowheads, **D,E**). **(F)** Quantification of basal protrusion phenotype by Kappa curvature analysis in ImageJ reveals loss of protrusions after IWR1 treatment (REF; see Methods). Green in figure 2B is autofluorescence. Blue fluorescence is Hoechst-stained nuclei. Scale bars = 5µm.

To determine whether canonical Wnt signaling was required to maintain the invasive mesenchymal character of new nephron cells, we treated gentamicin-injured zebrafish with the canonical Wnt inhibitor IWR1 at 7 days post injury (dpi) and imaged the basal surfaces of new nephron cells at 8 dpi for invasive basal protrusions. Twenty-four hour IWR1 treatment eliminated basal protrusions (figure 2D,E). The average basal surface curvature of multiple new nephrons (n=7) measured using ImageJ Kappa (see materials and methods, ^30^) showed a significant reduction in basal protrusions (p < 0.0014) in IWR1 treated fish (figure 2F), indicating a requirement for canonical signaling for invasive activity.

### Wnt ligands *wnt9b* and *wnt4* are required for new nephron tubule formation

*wnt9a* and *wnt9b* are both induced by acute kidney injury in the adult collecting system and distal tubules, the site of new nephron addition ^23^, while *wnt4* is expressed in new nephrons at the point of contact between new nephrons and target distal tubules (figure 2B), making these Wnts candidate signaling ligands for regulating new nephron formation.

Mutation in either *wnt4* ^31^ (*wnt4^uc55^, referred to as wnt4 -/- hereafter*) (figure 3A,C,E) or *wnt9b* ^32^ (*wnt9b^fb207^, referred to as wnt9b -/- hereafter*) (figure 3B,D,F) markedly reduced nephrogenesis following acute gentamicin injury while adult mesonephric nephron number, as determined by counting *nephrin+* glomeruli in uninjured adult kidneys, was not affected by either mutation (Supplemental figure 2A,B). The results show that both *wnt9b* and *wnt4* are not required for mesonephric kidney development but are essential for new adult nephron formation following injury. In 7 dpi *wnt4 -/-* mutant kidneys, new nephron aggregates that did form remained small and confocal sections of image stacks showed no invasive basal protrusions or proliferative tubule outgrowth (figure 3H) compared with wildtype (figure 3G). In addition, *wnt4* mutant new nephrons did not form a tubule lumen (figure 3H). Acute injury-induced *wnt9b* expression was not affected by the mutation in *wnt4* (Supplemental figure 3).

**Figure 3.**
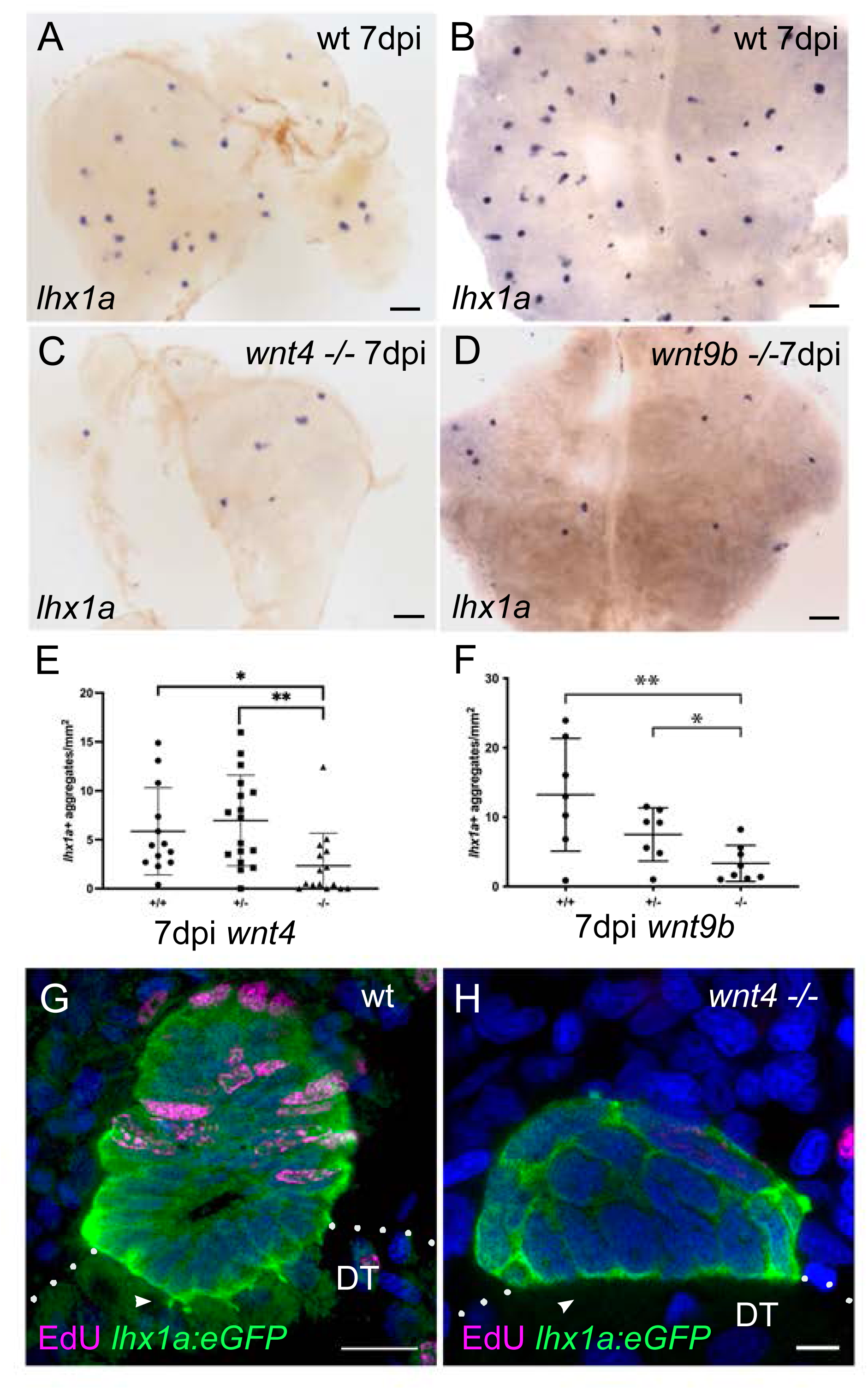
*wnt9b* and *wnt4* are required for new nephron formation and invasive cell behavior. Wildtype adult kidney **(A, B)** show a robust regenerative response to gentamicin acute injury 7 days post injection. Homozygous mutation in *wnt4* **(C)** or *wnt9b* **(D)** significantly reduces new nephron formation after acute gentamicin injury. Scale bars in A-D = 0.2mm. Quantification of *lhx1a+* new nephron aggregates in *wnt4* **(E)** and *wnt9b* (**F)** heterozygous and homozygous mutants reveals a significant reduction in nephrogenesis. New nephron aggregates in *wnt4* homozygous mutants show lack of cell polarization, failure of lumen formation, and absence of basal cell protrusions compared to wildtype (**G,H)**. Blue fluorescence is Hoechst-stained nuclei. Scale bar = 10 µm in G; 5 µm in H.

### Src kinase and Rac1 signaling are required for basal protrusion formation

Cellular protrusions, including invadopodia, are often initiated by localized Src family kinase activity ^33^. To determine whether Src kinases were involved in tubule interconnection, we treated gentamicin injured zebrafish at 7 dpi with the Src inhibitor PP2 and assessed basal surface curvature in confocal image projections. Twenty-four hour treatment with PP2 at 7-8 dpi eliminated basal protrusions on new nephron basal surfaces, while the negative control PP3 had little effect (figure 4A-C). Assessment of new nephron average basal surface curvature in PP2 treated using ImageJ Kappa revealed a significant inhibition of protrusive activity (n=13, p< 5 E-05) (figure 4C). Basal protrusions have also been shown to require the GTPase rac1 to direct actin-based protrusive activity REF. Involvement of rac1 was tested using the inhibitor EHT1864 ^34^. EHT1864 treatment disrupted the organization of actin filaments in new nephron basal protrusions (figure 4D, E). The normally central core of actin filaments in protrusions (figure 4D,D’) was dispersed to the plasma membrane and basal protrusions had a swollen, bleb-like appearance (figure 4E,E’). Basal protrusion cross sectional area of EHT1864 treated fish was significantly increased (figure 4F). These results reveal that invasive cell behavior associated with tubule interconnection requires Src kinases and rac1 activity.

**Figure 4.**
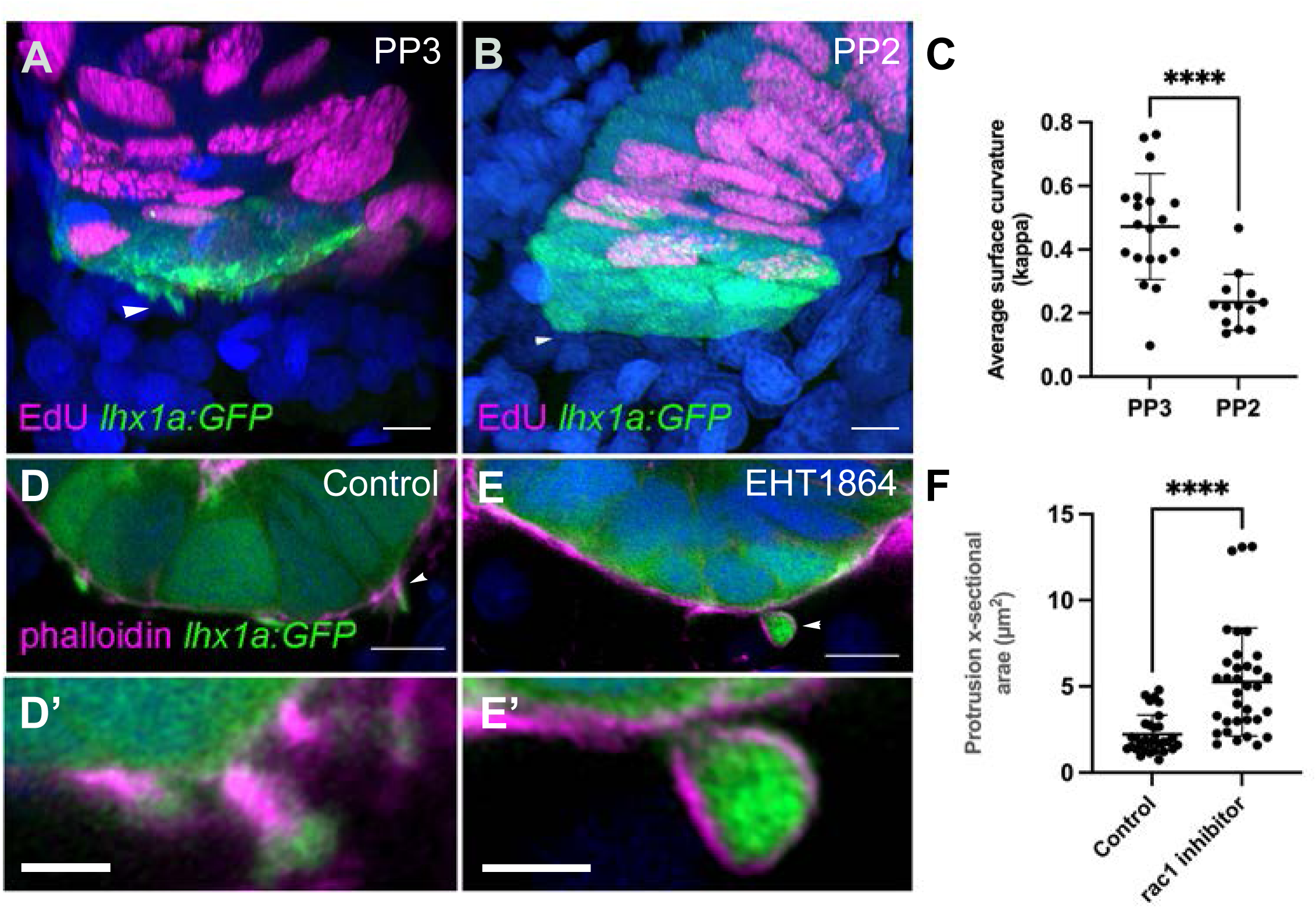
Src kinases and rac1 function in basal protrusion formation. Control (PP3) treated kidney **(A)** new nephrons show invasive basal protrusions (arrowhead) that are missing in Src family inhibitor (PP2; **B)** treated kidneys. Quantification of surface curvature **(C)** revealed PP2 caused a significant loss of basal protrusions. Basal protrusions in control new nephrons **(D, D’)** are formed with a central actin core and maintain a narrow, invasive morphology. rac1 inhibition with EHT1864 (**E, E’)** caused mislocalization of actin to basal protrusion cell membranes, distension of protrusions, and a “blebbing” morphology. EHT1864-induced distension of basal protrusions quantified **(F)** by protrusion cross-sectional area. Blue fluorescence is Hoechst-stained nuclei. Scale bars in A,B,D,E = 5µm; D’,E’ = 2µm.

### *frizzled 9b* mediates a non-canonical Wnt signaling pathway in tubule fusion

Frizzled 9b (*fzd9b*) is a Wnt receptor expressed in adult zebrafish kidney progenitor cells and in newly forming adult nephrons^23^ (figure 5A,B). Mutation in *fzd9b* causes a roughly 50% reduction in mesonephric nephron number and a roughly 50% reduction in new nephron induction after acute kidney injury ^23^. To determine if *fzd9b* was responsible for canonical or non-canonical Wnt signaling at the point of tubule interconnection, we examined canonical Wnt reporter activity and endogenous canonical Wnt target gene expression in *fzd9b* homozygous mutant kidneys at 7 dpi. Unexpectedly, mutation in *fzd9b* resulted in expanded expression of the canonical Wnt reporter *Tg(tcflef:dGFP)* (figure 5C,D) and the endogenous canonical Wnt target gene *wnt4* (figure 5E,F). Similarly, the expression domains of canonical Wnt target transcription factors *lef1*, *lhx1a*, and Wnt-induced negative feedback inhibitors *notum1*, and *wif1* were expanded in *fzd9b* mutants (Supplemental figure 4), suggesting that *fzd9b* normally acts to antagonize canonical Wnt signaling. We have shown that canonical Wnt signaling is required for cell proliferation in new tubule aggregates ^23^. Consistent with an antagonistic function for *fzd9b*, cells at the point of interconnection in new tubules that do not normally proliferate (figure 1H, 6A) exhibit active cell proliferation in the *fzd9b* mutant (figure 6B). Taken together, the data indicate that *fzd9b* does not mediate canonical signaling in the context of tubule interconnections but rather antagonizes canonical Wnt signaling, most likely via a non-canonical Wnt pathway. In support of this idea, we found that additional non-canonical Wnt signaling components, *prickle2b* and *ptk7,* were expressed in newly forming kidney tubules (Supplemental figure 5).

**Figure 5.**
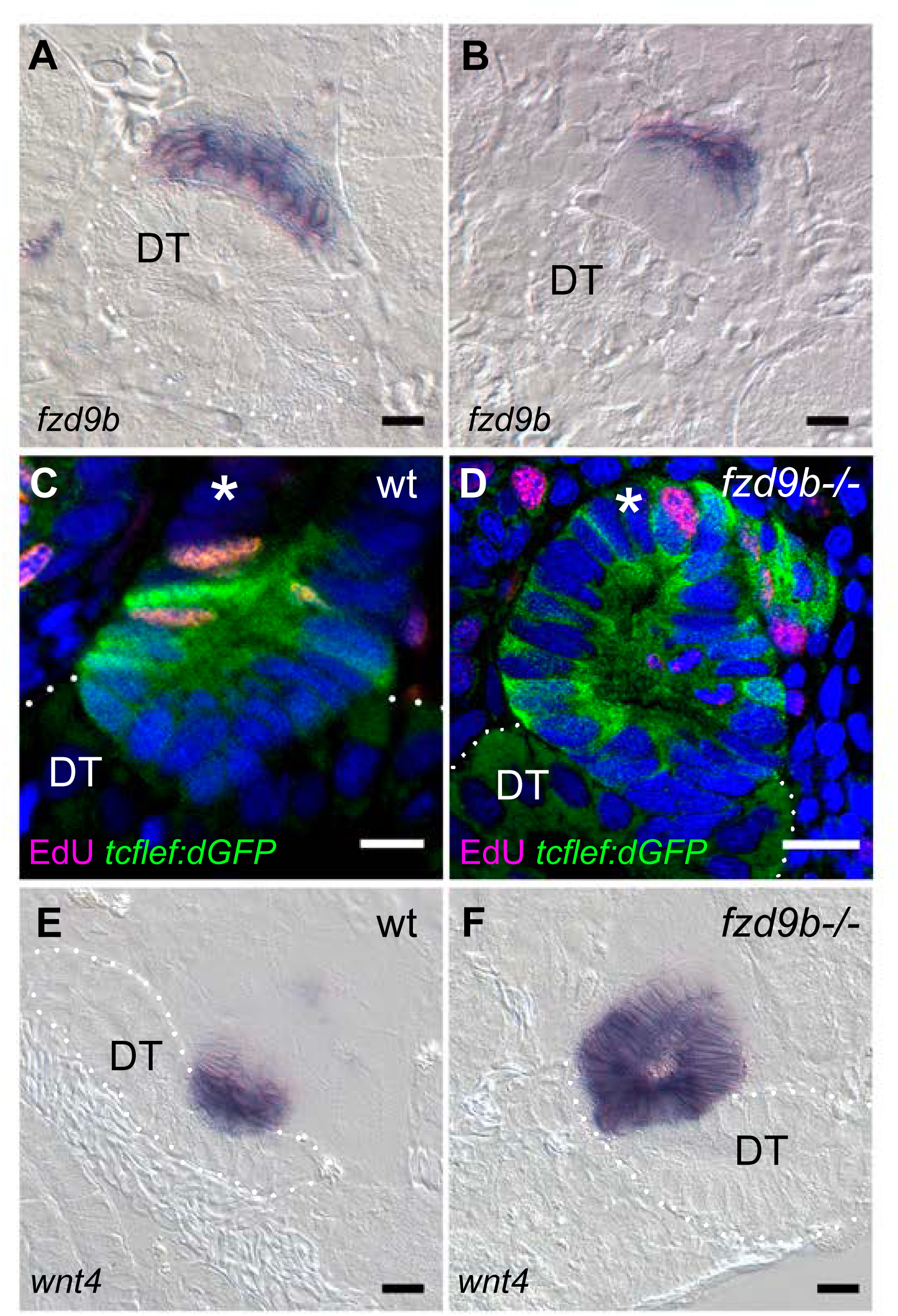
*frizzled9b* antagonizes canonical Wnt signaling. The Wnt receptor *fzd9b* is expressed in new nephrons adjacent to the distal tubule **(A**; DT = distal tubule) and in some cases, several cell diameters away from the junction surface (**B**). Expression of the canonical Wnt reporter *Tg(tcflef:dGFP)* (**C**) is restricted to new nephron cells adjacent to the distal tubule (DT) and not expressed in cells several cell diameters away (asterisk). Mutation in *fzd9b* results in expanded expression of *Tg(tcflef:dGFP)* where the entire new nephron can show GFP expression **(D**, asterisk). (**E**) *wnt4,* an endogenous canonical Wnt target gene, is expressed in a domain restricted to new nephron cells adjacent to the distal tubule (DT). Mutation in *fzd9b* results in significantly expanded *wnt4* expression **(F**). Blue fluorescence is Hoechst-stained nuclei. Scale bars in A,B,E,F = 20 µm; C,D = 10µm.

**Figure 6.**
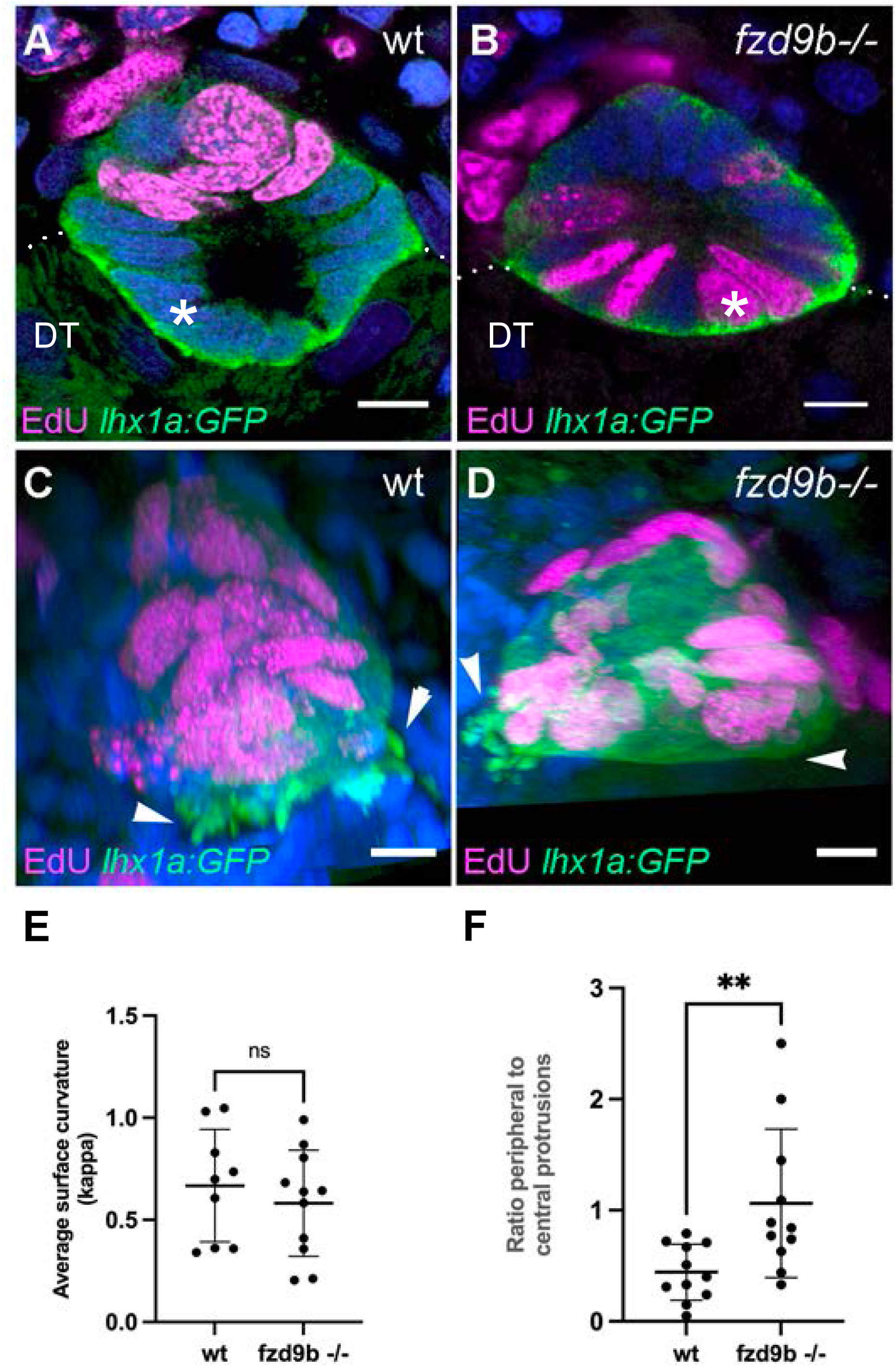
*frizzled9b* is required to limit cell proliferation and direct basal protrusions to generate orthogonal tubule interconnections. Basal new nephron invading cells are non-proliferating in wildtype **(A)** but show ectopic proliferation in *fzd9b* mutants **(B)**. Basal protrusions are directed orthogonally at the target distal tubule in wildtype **(C)** but show a lateral displacement in fzd9b mutants **(D**; arrows). Overall surface curvature of the invading surface is not changed in the *fzd9b* mutant (**E**; arrowheads) however the basal protrusions are mislocalized laterally (**F**; arrowheads). Blue fluorescence is Hoechst-stained nuclei. Scale bars = 5µm.

Non-canonical Wnt signaling pathways drive planar cell polarity and modulate the cytoskeleton to regulate cell shape and orient collective cell movements during tissue morphogenesis and gastrulation ^35^. During interconnection, new tubules manifest several cellular features associated with non-canonical Wnt signaling including apical constriction of new tubule epithelial cells (figure 1A,C) and cell protrusions associated with cell movement that precedes lumen fusion (figure 1G) ^36^. In wildtype kidneys, basal protrusions in new tubule aggregates are distributed across the tubule cell aggregate basal surface and oriented parallel to the invading lumen (figure 6C) to ultimately create an orthogonal junction between two epithelial tubes (figure 1B). Mutation in *fzd9b* disrupted this organization and mispolarized cell protrusions laterally such that they were misdirected away from the target distal tubule lumen (figure 6D; supplemental movie 4, compare with figure 1G; supplemental movie 3). Overall protrusive activity was not significantly impaired (figure 6E) however the distribution of protrusions was shifted to the periphery of tubule basal surfaces (figure 6F). Apical constriction of new tubule epithelial cells was also disrupted in the *fzd9b* mutant. Comparison of the apical cell surface in wildtype (figure 7A) and *fzd9b* mutant tubules (figure 7B) prior to interconnection revealed that the apical area in *fzd9b* mutants was increased more than 6-fold (figure 7D) leading to cystic expansion of the new tubule lumen abutting on the distal tubule. New nephron lumen growth was directed laterally in the *fzd9b* mutant (figure 7B, supplemental movie 5) instead of toward the target distal tubule lumen. Non-canonical Wnt signaling drives apical constriction via rho and rho kinase, leading to myosin light chain phosphorylation and contraction of apical actomyosin to generate cell shape change ^37^. We found that treating 7 dpi fish with the ROCK inhibitor Y-27632 for 24 hours phenocopied the *fzd9b* mutant and caused a greater than 3-fold increase in apical cell surface area while the rac1 inhibitor EHT1826 had no effect on apical cell surface area (figure 7C, D). Invading cells of *fzd9b* mutants exhibited neither apical nor basal constriction ( (figure 7E,F; compare figure 7F to figure 1E,F) indicating a loss of dynamic cell shape change in the mutant. Final lumenal connections in wildtype tubules were orthogonal and patent (figure 7G) whereas in *fzd9b* mutants, connections were constricted when they did occur and lumens were circuitous, often migrating around a target distal tubule before making a connection (figure 7H, supplemental movie 6).

**Figure 7.**
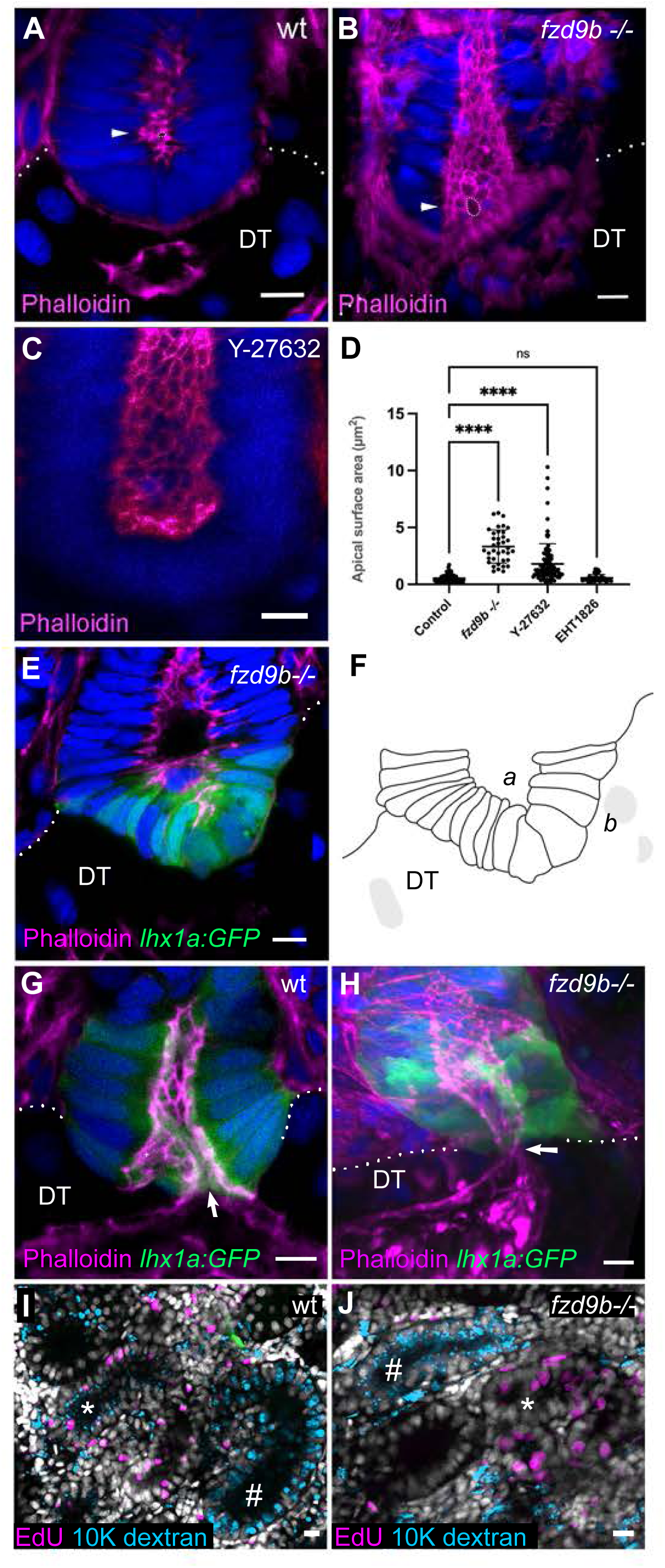
*frizzled9b*-dependent apical constriction guides orthogonal interconnection of tubules and functional fluid filtration. New nephron lumens are narrow and composed of apically constricted cells with small apical surfaces (**A**; arrow, outline). *fzd9b* mutants lose apical constriction, show distended lumens, and fail to make orthogonal interconnections (**B**; arrow, outline). (**C**) Apical constriction requires Rho kinase signaling. The ROCK inhibitor Y-27632 phenocopies the *fzd9b* mutant phenotype. (**D**) Quantification of apical surface area in *fzd9b* mutant, ROCK/Y-27632 inhibited, and rac1 / EHT1826 inhibited new nephrons. (**E**) *fzd9b* mutant new nephron cells fail to undergo cell shape change from apical to basal constriction (see figure 1) and instead maintain a primarily rectangular shape with roughly equivalent apical (*a*) and basal (*b*) surface areas (**E, F**). Compared to wildtype tubule interconnections (**G**; arrow), *fzd9b* mutant interconnections (**H**) are delayed and narrow (arrow). (**I**) In wildtype kidneys at 14 dpi, intraperitoneally injected 10K fluorescent dextran marks both proliferating, EdU+ new nephrons (*) as well as non-proliferating pre-existing nephrons (#). (J) In *fzd9b* mutants, pre-existing nephrons take up injected dextran (#) however, EdU+ new nephrons show reduced frequency of dextran uptake. Blue fluorescence is Hoechst-stained nuclei. Scale bars = 5µm.

To functionally assess tubule continuity, we performed intraperitoneal injections of 10kDa fluorescent dextran in adult fish 12 days after gentamicin injury and identified newly formed tubules using EdU incorporation. In this protocol, only newly forming tubules are marked by EdU since EdU is injected at 12 and 14 dpi, well after older tubules have responded to injury but during the period of new nephron growth. Dextran uptake was readily apparent in older, pre-existing tubules in both wildtype and *fzd9b* mutant kidneys (figure 7I,J; non-proliferating tubules marked with #). In wildtype, EdU+ new tubules were also positive for dextran (figure 7I; 10/12 tubules; 83%) indicating fluid flow and successful connections had been established. In contrast, *fzd9b* mutant tubules showed minimal dextran uptake (figure 7J; 6/17 tubules; 29%), consistent with reduced or absent lumenal perfusion due to constricted or failed tubule connections. In the kidney, tubule obstruction results in tubule swelling and distension due to blocked outflow and lumenal pressure ^38^. Consistent with failed interconnection and constricted flow, we observed large, distended tubule lumens in *fzd9b* mutant kidneys but not wildtype (Supplemental figure 6). Overall, these findings suggest that fzd9b enables tubule invasion and orthogonal lumen interconnection by transducing non-canonical Wnt signaling that controls both spatially restricted formation of invasive protrusions and the concerted apical to basal cell constriction required for epithelial tubule integration. Our studies also reveal a novel feature of morphogenetic cell signaling, where mutually inhibitory canonical and non-canonical Wnt pathways are sequentially engaged in the same cells to drive programs of proliferation versus invasion.

## Discussion

Our analysis of epithelial tubule fusion in the regenerating zebrafish kidney revealed that a complex interplay of Wnt signaling systems mediate the formation of patent, orthogonal tubule interconnections and ensure fluid flow in newly engrafted kidney nephrons. We found that both canonical and non-canonical Wnt signaling pathways operate simultaneously in the same cells to orient and drive invasive cell behavior and permit cell shape changes that facilitate tubule interconnection.

### Canonical Wnt signaling in tubule interconnection

Canonical Wnt signaling was required to induce a mesenchymal, invasive cell phenotype at the new nephron / distal tubule point of interconnection (figure 8A). This invasive phenotype was supported by expression of several genes typically associated with cancer metastasis (Supplemental figure 1). Generation of invasive protrusions was blocked by the canonical Wnt inhibitor IWR1 and mutant analysis revealed that autocrine *wnt4* signaling was required for basal protrusion formation. This pathway also required Src kinase activity. In human pathology, Wnt4 is known to be a driver of human invasive lobular carcinoma and colorectal cancer cell proliferation, cancers associated with invasive cell behavior or epithelial mesenchymal transformation ^39,40^. Src family kinase activity is also observed in the context of multiple other Wnt-dependent, invasive tumors ^41,42^ and is a prominent feature of invadopodia, invasive structures characterized by Mmp-14-and Tks5-expression ^43,44^. Like the zebrafish kidney, Wnt4 signaling in mammary carcinoma and thymic epithelial tumors is largely autocrine ^45,46^ and, in some cases, occurs independently of typical pathways for Wnt secretion ^47^.

**Figure 8.**
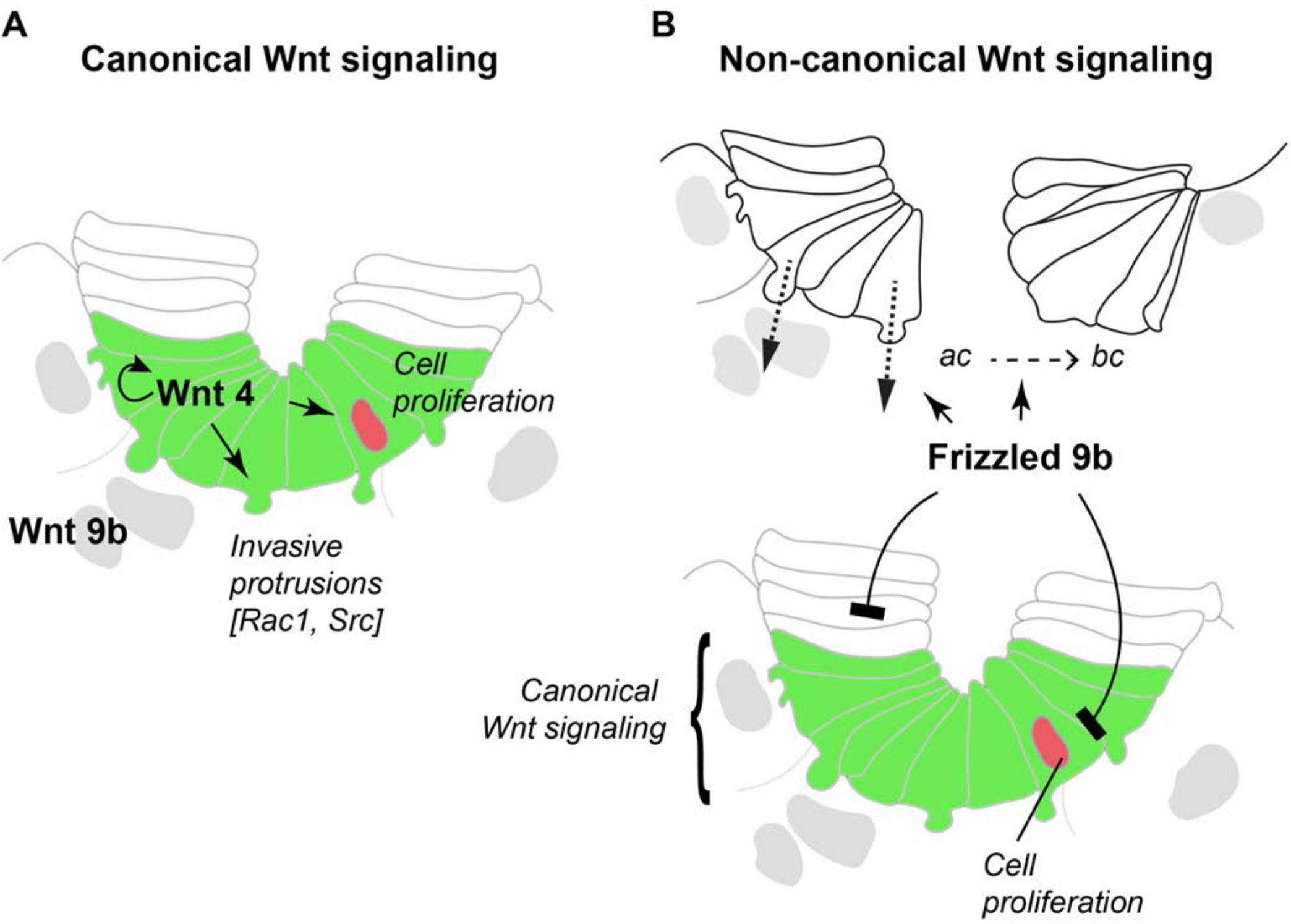
Multiple Wnt signaling pathways mediate tubule interconnection. **(A)** *wnt9b* expressed in the distal tubules (DT) is required for kidney regeneration and development of new nephrons. *wnt4* is induced at the junction of new and existing nephrons and appears to act in an autocrine fashion to promote cell proliferation, induce formation of basal protrusions, and polarize new nephron epithelial cells. Invasive basal protrusions require canonical Wnt signaling as well as Src kinase and Rac1 GTPase activity. **(B)** Non-canonical Wnt signaling mediated by Frizzled 9b activity mediates cell shape change (apical to basal constriction; ac --> bc) in a rho kinase dependent fashion and restricts canonical Wnt signaling to a narrow band of cells abutting existing distal tubules (green). Frizzled 9b activity also limits cell proliferation and positions basal protrusions orthogonally to distal tubules to promote patent tubule interconnections.

Invasive basal protrusions in new nephrons also required Rac1 activity to maintain a dense core actin organization. Rac1 inhibition caused protrusions to become cytoplasmic blebs which may also support cell migration ^48^. Rac1-dependent morphogenetic processes driven by basal protrusions are commonly observed in developmental contexts including Drosophila germband extension, ^49^, mammalian endothelial cell rearrangement during angiogenesis ^50^, and epithelial intercalation of the C. elegans embryonic epidermis ^51^. Our results on new tubule invasion in the adult zebrafish kidney present a useful model to study invasive epithelial morphogenesis in the context of a functioning adult organ.

Invasive cells do not proliferate in the regenerating zebrafish kidney. Cell cycle arrest linked to invasive behavior has been noted in both developmental contexts and in cancer metastasis ^52–54^. Studies in C.elegans show that cell invasion is a differentiated cell state that requires G1 arrest and is under genetic control by the transcription factor NHR-67/TLX (nuclear hormone receptor family) and HDAC-mediated changes in gene expression ^54^. In our study, suppression of proliferation in invading cells required a *fzd9b* regulated pathway since mutation in *fzd9b* relieved this block on proliferation. We also found that *fzd9b* signaling suppresses canonical Wnt signaling and Wnt target gene expression, suggesting *fzd9b* in new nephrons activates an antagonistic, non-canonical Wnt pathway that patterns the nephron, restricting canonical Wnt signaling to a sharp band of cells just adjacent to the target distal tubule. This activity of *fzd9b* stands in contrast to its role in hematopoietic stem cell emergence where a *wnt9b* - *fzd9b* interaction drives canonical Wnt signaling and activation of the LRP6/ß-catenin pathway^55^.

We have shown previously that canonical Wnt signaling promotes new nephron proliferative outgrowth ^23^. Our results here show that *wnt4* expression in the new nephron is required for nephron cell proliferation, epithelial cell polarization and generation of basal protrusions (see above). Since *wnt9b* continues to be expressed (locally in the target distal tubule) in *wnt4* mutants and is not sufficient to promote proliferation in new nephrons, *wnt4* may be the primary Wnt ligand required for cell proliferation in this context. Our results are consistent with data from mouse kidney development where Wnt9b in the ureteric epithelium is required to induce Wnt4 in nephron progenitors which subsequently drives nephron epithelialization ^27,56^. While additional allelic combinations of *wnt4* and *fzd9b* will help elucidate the exact relationship between *fzd9b* and *wnt4* signaling, our results point to a *fzd9b* pathway that maintains cell quiescence, possibly similar to non-canonical Wnt pathways that maintain quiescence in neural stem cells ^57^, mouse muscle satellite cells ^58^, and Xenopus kidney ^59^.

### Non-canonical *fzd9b* signaling mediates morphogenetic transitions required for tubule interconnection

In addition to its function in limiting cell proliferation and patterning canonical Wnt target gene expression in new nephrons, *fzd9b* was required for apical constriction of new nephron epithelial cells at the fusion interface and a transition to basal constriction as interconnection progressed. Apical and basal constriction are universal developmental mechanisms in epithelial sheets undergoing shape change or invagination as occurs in gastrulation ^60,61^, optic cup formation ^62^, and brain morphogenesis ^63^. A recent report on Xenopus neurulation highlights a role for a Wnt4 / ephrinB2 non-canonical Wnt pathway that activates rho GTPase dependent acto- myosin contraction to effect apical constriction in the neural plate ^64^. Our data show that a non-canonical Wnt pathway involving *fzd9b* signaling can similarly act in tubular epithelia to activate rho pathway-dependent apical constriction. Cells abutting the target distal tubule in *fzd9b* mutant kidneys also showed a mislocalization of basal protrusions to the periphery of the connection interface. This result may reflect an overall failed polarization of cells in the *fzd9b* mutant. The result of mis-localized protrusions and failed cell shape change was that new nephrons did not fully penetrate the target distal tubule, and no orthogonal connections were made. Instead, the growing lumen was displaced laterally and, in some cases, projected around the target distal tubule to form a partially patent connection.

Beyond the basic knowledge of how epithelial cells and tubules interconnect, our data has practical implications for utilizing stem cell-derived kidney organoid nephrons in tissue replacement and kidney augmentation therapies ^15,65^. Attempts to engraft and fully integrate organoid derived nephrons in an adult mouse kidney have been challenging due to the lack of patent nephron interconnections with the host tubular architecture ^15^. Connections to the host and establishing functional nephron “plumbing” will be essential to generate nephron fluid flow and blood filtration. Our results provide new knowledge of cellular activities and signaling systems that may be engineered at an organoid / host tubule interface to promote tubule interconnection and function of engrafted nephrons.

### Limitations of the study

We have defined two ligands, Wnt9b and Wnt4, and one receptor, Frizzled9b, as essential for new nephron formation and described their role in interconnection of new nephrons to existing tubules. However, which ligands bind to *fzd9b* or other Wnt receptors has not been fully defined. Loss of function of either the Wnt9b or Wnt4 ligands or the *fzd9b* receptor has some opposite effects on the patterning of new nephrons (activation vs. repression of canonical Wnt target gene expression) indicating that *fzd9b* is unlikely to mediate canonical Wnt signaling and induction of basal protrusions in this context. We have not mutated Wnt co-receptors (*ror2, ptk7*) that may play a defining role in directing Wnt signaling to the canonical or non-canonical pathways. Although there is much remaining to do, this work defines an important framework to understand the complex interplay of the competing Wnt signaling pathways that drive tubule interconnection.

## Methods

### Fish care, injury and drug treatments

Wild-type TuAB zebrafish were maintained according to established protocols ^66^. Each experiment was performed with age-matched siblings reared together to minimize background genetic variation when possible. All adult experiments were performed with fish between 6-18 months of age. Acute kidney injury was induced by intraperitoneal injection of gentamicin as previously described ^24^. Fish weighed between 0.25g and 1.0g. Gentamicin (Sigma Aldrich) diluted in PBS was injected at 80mg/Kg. Drug treatments were performed by keeping fish in either 5uM (IWR1, IWP2) or 10uM (PP2, PP3) or 100uM (Y-27632) (Tocris) or DMSO (Sigma) dissolved in system water starting at 7dpi until harvesting kidneys at 8dpi, except for EHT1864 which was injected i.p. at a dose of 50mg/Kg. Water changes were performed every other day by replacing half of the volume with fresh drug treated water. For short term proliferation studies, 20ul of 05mg/ml EdU (Invitrogen) dissolved in HBSS (Sigma Aldrich) was delivered by intraperitoneal injection 2 hours before euthanization. For longer term interconnection studies, EdU was injected at 6dpi, drug treatments were done at 9dpi, and kidneys were harvested at 10dpi. For dextran filtration experiments, EdU was injected at 10 and 12dpi, 20 µl 10KD rhodamine dextran (1mg/mL in PBS) (Thermo Fisher) was injected at 12dpi and kidneys were harvested at 14dpi.

### *in situ* hybridization and immunofluorescence

Whole-mount single *in situ* hybridization was performed as described ^68^ with some modifications ^24^. Briefly, fish were fixed with the head and internal organs removed, leaving the kidneys attached to the dorsal body wall, overnight with rocking in 4% paraformaldehyde (Electron Microscopy Sciences). After washing 5 times with PBST (PBS with 0.1% Tween-20), fixed kidneys were removed from the body using forceps and permeabilized with proteinase K (10ug/ml Roche) in PBST for 1 hr with rocking, postfixed in 4%PFA overnight and washed 5 times with PBST. Probes for *mmp14a*, *mmp14b*, *tks5*, *cdh11*, *cjun* and *notum1a* were cloned from 2dpf TuAB cDNA using the indicated primers using a hot start PCR with Phusion polymerase (NEB) and cloned in to PCR II Blunt TOPO vector (Thermo Fisher). These and *lhx1a, fzd9b, lef1, wnt4, wnt9b, nephrin, id1, prickle1b, ptk7, notum1* and *wif1* ^69,70^ REFs were synthesized using DIG RNA labeling mix (Sigma Aldrich). After staining, kidneys were fixed with 4% PFA, cleared with dimethylformamide, depigmented with hydrogen peroxide, transferred into PBS:glycerol (1:1) and imaged on a Leica MZ12 microscope equipped with a Spot Image digital camera. Dehydrated kidneys were embedded in JB-4 plastic resin (Polysciences) and then sectioned to a thickness of 7 μm using a LEICA 2065 rotary microtome and mounted using Permount (Fisher Scientific) with #1 coverslip (Electron Microscopy Sciences). Sections were imaged on a Nikon E800 microscope equipped with Plan Apo 60xA/1.4 oil objective with a Spot Insight CCD digital camera.

Quantification of *lhx1a+* aggregates was performed in a blinded manner using ImageJ. Briefly, the number of aggregates in a single 5x image taken of the widest section of each kidney was divided by the total kidney area measured in ImageJ using the freehand tool to calculate aggregates per mm^2^. Pictures were relabeled before counting such that the person doing the analysis had no knowledge of treatment conditions for each sample.

For immunostaining, the initial fixation step was 3 hours instead of overnight but otherwise the same. Kidneys were stained for GFP (chick anti-GFP, 1:5000 (Abcam ab13970) and goat anti-chick Alexa Fluor 488, 1:3000 (Thermo Fisher)). After antibody staining, kidneys were treated for EdU detection with the Click-iT Edu kit (Invitrogen) according to manufacturer’s instructions with a 1 hr incubation and stained with Hoechst 33342 (1:2000, Invitrogen) overnight to detect nuclei. For double in situ hybridization and antibody staining, probe was detected using the Tyramide Super Boost kit (Invitrogen) followed by antibody staining for GFP (A11122 rabbit anti-GFP, 1:500, (Invitrogen), goat anti-rabbit Alexa Fluor 488 1:3000 (Thermo Fisher)). Phalloidin labeled with Alexa Fluor 594 (1:400, Thermo Fisher) was added to overnight Hoechst staining to visualize F-actin. Stained kidneys were mounted in PBS:glycerol (1:1) on a 1 coverslip high bridge slide using #1.5 coverslips glued down with Krazy glue (WB Mason) for imaging.

Laminin (L9393, 1:30 (Sigma), goat anti-rabbit Alexa Fluor 546 1:3000 (Thermo Fisher)) and GFP staining were performed on 20um cryosections. Briefly, kidneys were infiltrated with 30% sucrose/PBS overnight, embedded in OCT medium (TIssueTek) and sectioned on a Leica CM1860UV cryostat, stained in a humidified chamber then mounted with Vectashield medium (Vectashield) under a #1.5 coverslip for imaging.

Confocal imaging was performed on a Zeiss LSM980 point scanning confocal microscope with Airyscan 2 (Carl Zeiss Microscopy) equipped with a Zeiss Axio Examiner Z1 upright microscope stand (ref:409000-9752-000) and a Plan/Apochromat 10x/0.45 (ref:420640-9900-000) or Plan/Apochromat 63x/1.4 oil (ref:420782-9900-799) objective controlled with Zen Blue software (Zen Pro 3.1). Z stacks taken with the 10x objective were acquired using Nyquist criterion mode. Z stack images at 63x were collected with the Super Resolution mode (SR) at either 1.7x or 3.4x zoom with optimal interval using the motorized scanning stage 130 x 85 PIEZO mounted on the Z-piezo stage insert WSB500 (Carl Zeiss Microscopy). Airyscan images were processed using the “auto” mode and saved in CZI format. 3D reconstructions of Z stacks were created using Imaris 9.6 Software.

Quantification of dextran filtration was done by two separate observers using both Imaris 3D reconstructions and stacks of slices in ImageJ and results were reconciled. Newly regenerated nephrons were identified by searching stacks for EdU+ tubules (identified by epithelial nuclear morphology and EdU+ nuclei) with the rhodamine dextran channel turned off, then counting the number of tubules which had dextran containing vesicles on the apical/luminal side of nuclei.

#### Cell membrane curvature analysis

Basal protrusions on new nephrons were quantified by measuring average basal surface curvature of *lhx1a:egfp+* new nephrons. Zeiss .czi files were converted to Imaris format (.ims) and free rotated to be orthogonal to the viewing axis and exported as an image sequence. The ImageJ Kappa plugin ^30^ was used to analyze individual slices every 2 µm of stack depth using tracings of new nephron bottom surfaces outlined by *lhx1a:egfp* fluorescence. Final measurements were exported to Excel and Prism to average the kappa measurements of individual new nephron basal surfaces.

#### Basal protrusion position analysis

Zeiss .czi files were converted to Imaris format (.ims) and free rotated to present an *en face* view of new nephron tubule basal surfaces. A region of interest was drawn, encompassing the entire basal surface then scaled to 90%. The ImageJ process “find maxima” (prominence 15.0) was used to generate a ratio of *lhx1a:egfp* fluorescence intensity maxima inside (central) vs. outside (peripheral) the ROI and quantified as a proxy for basal protrusion position on new nephron basal surfaces.

#### Apical surface area analysis

Apical surface area was measured in individual confocal slices of phalloidin stained new nephron confocal stacks. Phalloidin+ lumenal cell apical perimeters were traced and measured in ImageJ using the area measurement tool. Average apical surface area was computed using Excel and graphed using Prism.

## Acknowledgements

This research was supported by the National Institutes of Health (NIDDK 5UC2DK126021 to I.A.D., D.M., L.O.), NSF grant (IOS-1456737) to B.D., and NIGMS IDeA Award under P20GM103423, P20GM104318, and P30GM154610. Image collection, processing and analysis for this manuscript was performed with the assistance of the MDI Biological Laboratory Light Microscopy Facility (RRID:SCR_019166).

## Author Contributions

Conceptualization, C.N.K. and I.A.D.; Methodology, C.N.K., H.S., F.B., C.P.C., K.W.M., D.K.M., L.O. and I.A.D.; Investigation, C.N.K., W.G.B.S., C.A., O.A, A.W., R.U., S.M.H., M.A.,; Resources, B.W.D.; Writing-Original Draft, C.N.K. and I.A.D.; Writing – Reviewing and editing, all authors; Supervision, I.A.D.; Funding Acquisition, I.A.D, D.K.M., L.O., and B.W.D.

## Supplemental Figure Legends

**Supplemental figure 1.**
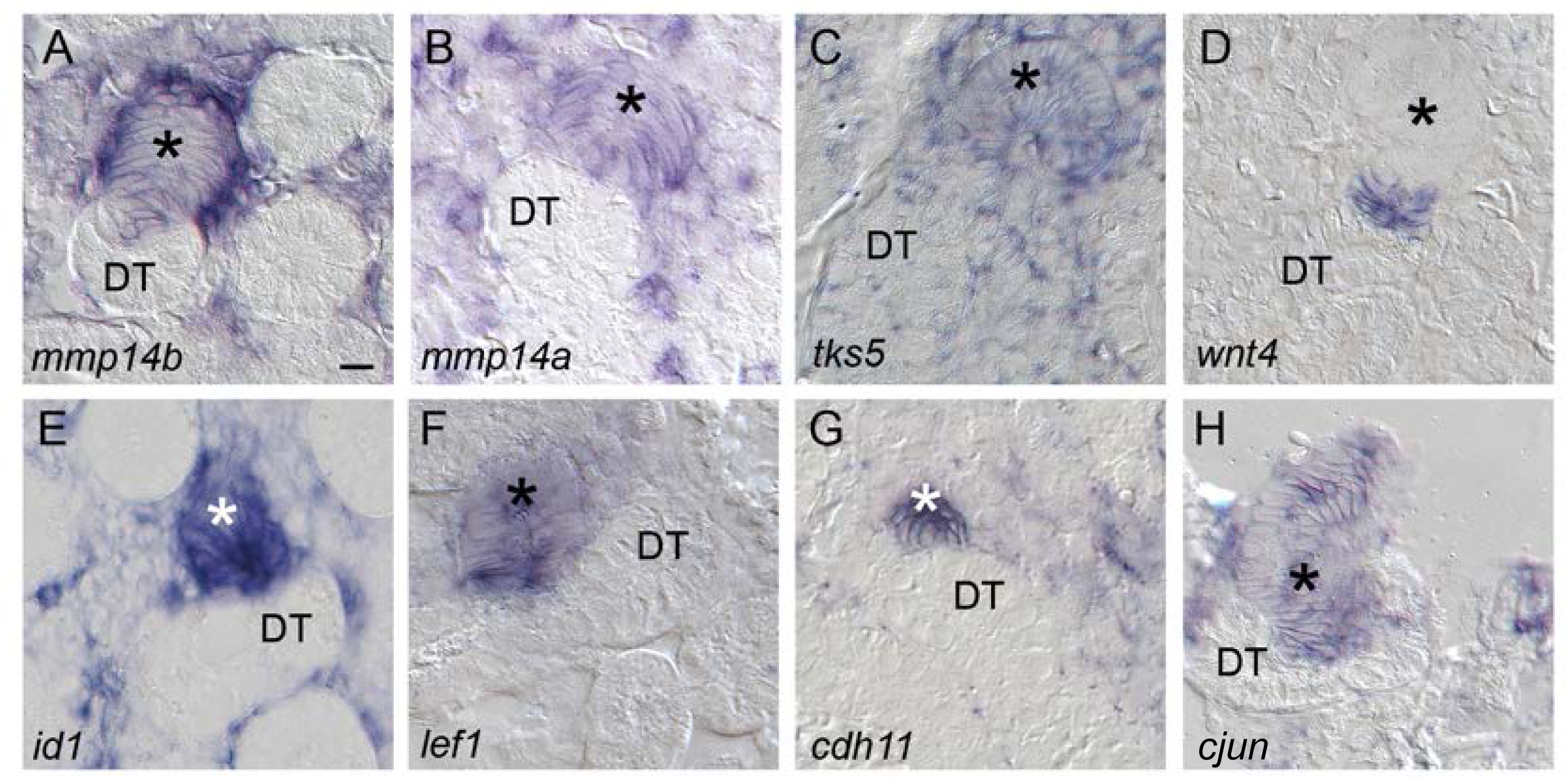
Expression of genes in connecting nephrons orthologous to human genes associated with invasive metastasis. Digoxigenin whole mount in situ hybridization, glycol methacrylate emedded and sectioned 7 dpi kidneys (* marks new nephron structures; DT = Distal Tubule; gene names noted in each panel)

**Supplemental figure 2.**
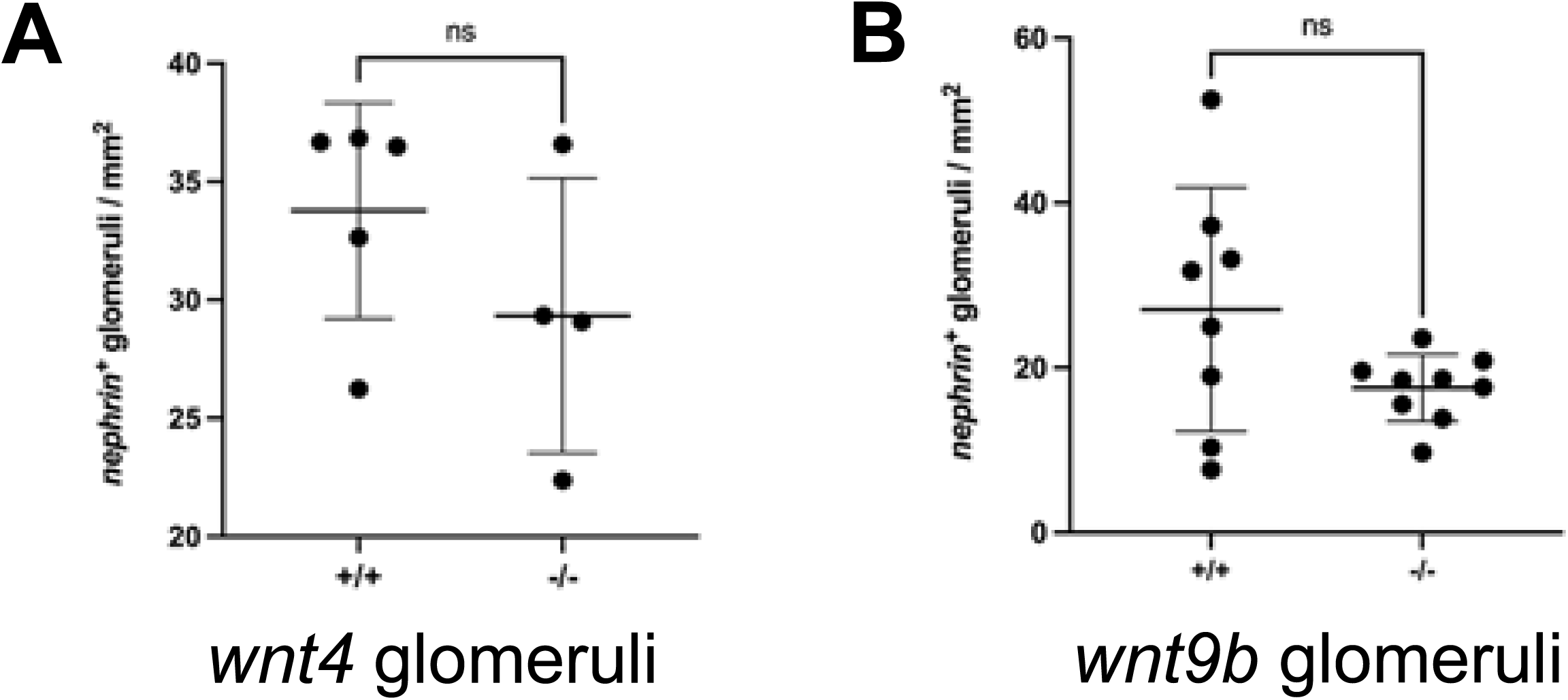
Quantification of nephron number by nephrin (glomeruli) whole mount in situ hybridization in *wnt4* and *wnt9b* mutants. **(A)** Comparison of nephron number between uninjured adult wildtype +/+ and homozygous *wnt4* mutant mesonephroi (-/-) shows no statistical difference. **(B)** Comparison of nephron number between uninjured adult wildtype (+/+ and homozygous *wnt9b* mutant mesonephroi (-/-) shows no statistical difference.

**Supplemental figure 3.**
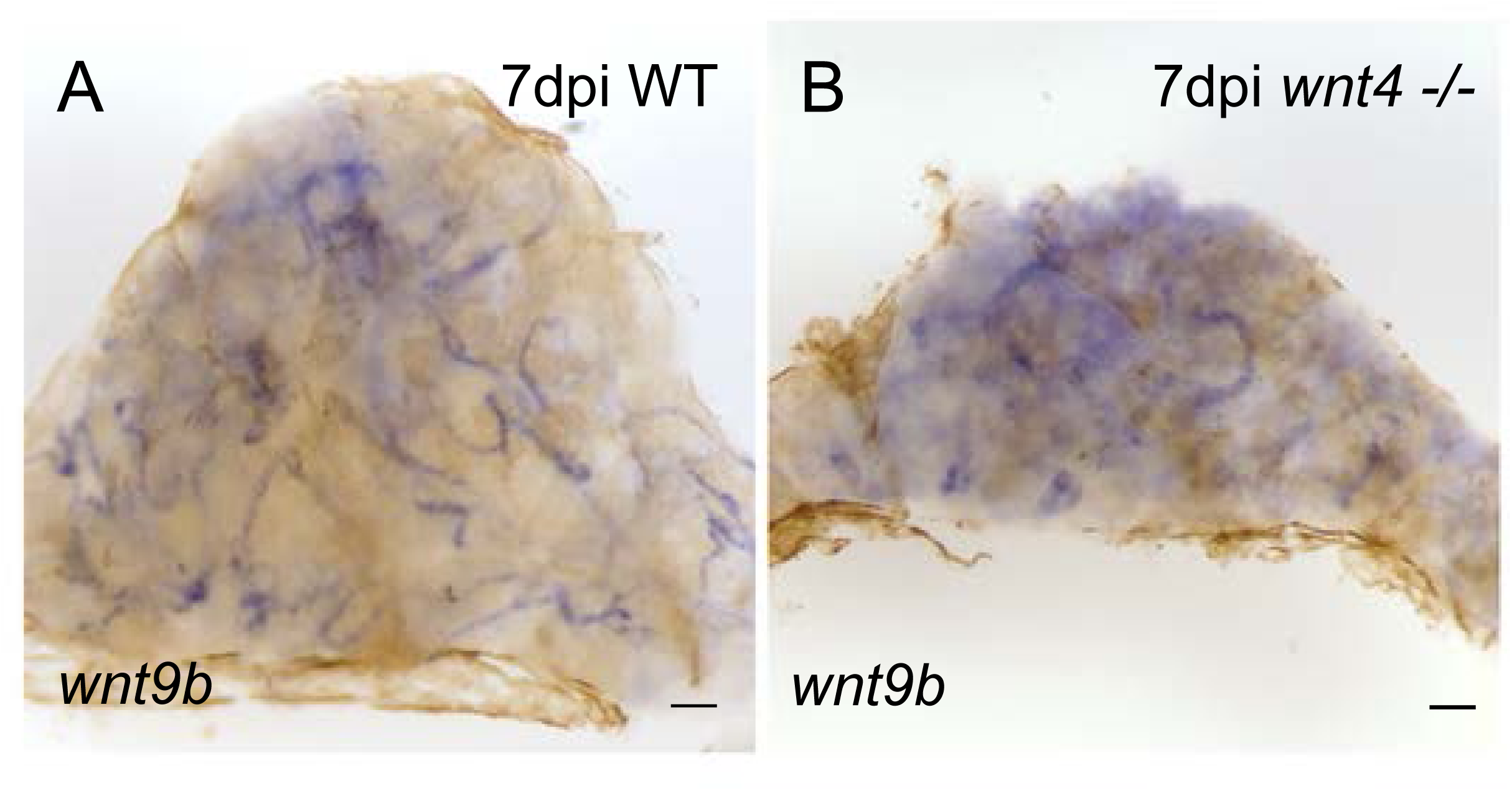
Distal tubule expression of *wnt9b* in 7 dpi adult kidney. **(A)** Whole mount in situ hybridization of *wnt9b* in distal tubules of wildtype kidney 7 days after acute gentamicin injury. **(B)** Whole mount in situ hybridization of *wnt9b* in distal tubules of *wnt4 -/-* kidney 7 days after acute gentamicin injury. Scale bars = 100µm.

**Supplemental figure 4.**
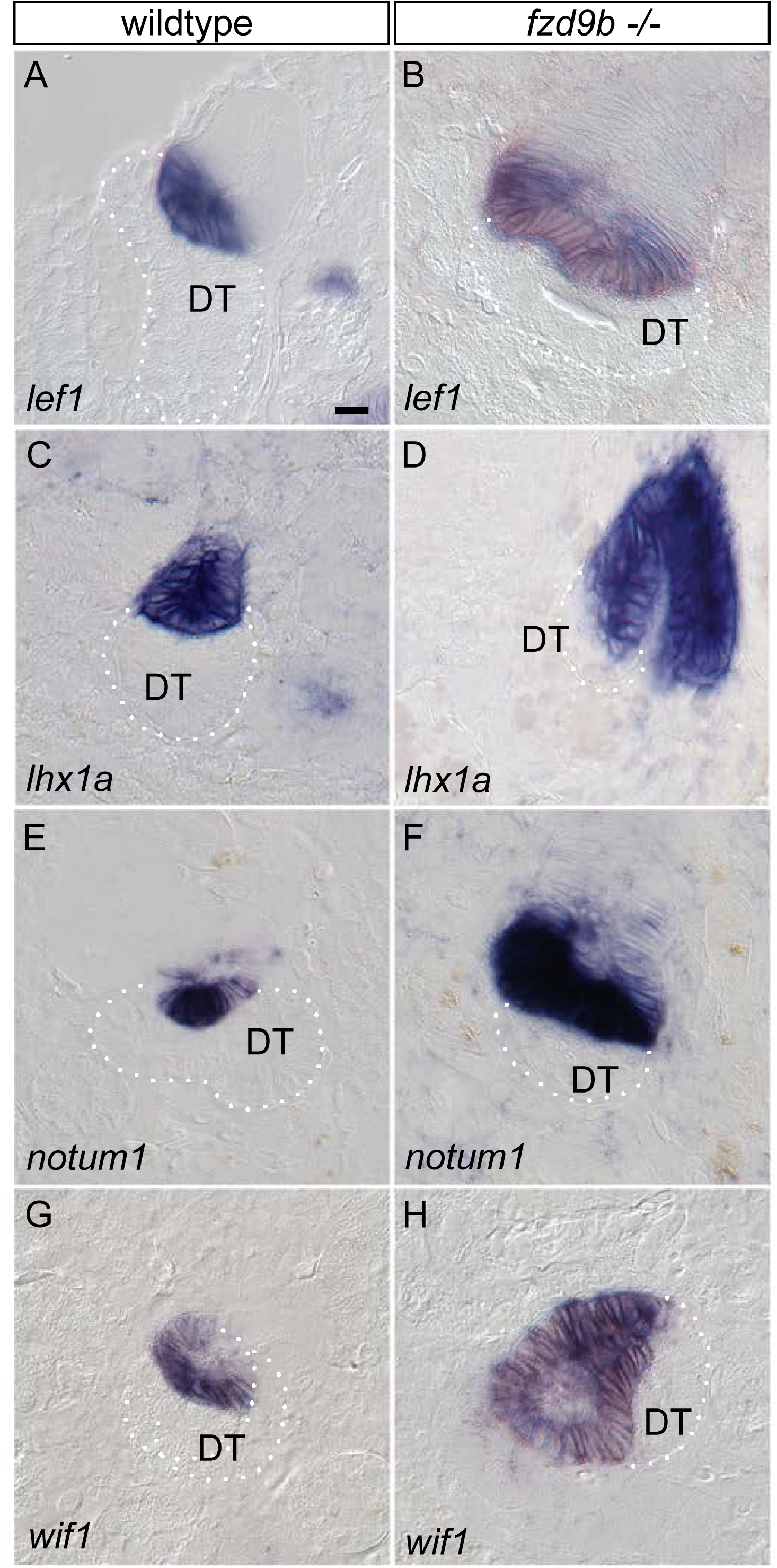
Expression of canonical Wnt signaling target genes in wildtype and *fzd9b* homozygous mutants. (A,B) *lef1*. **(C,D)** *lhx1a*. **(E,F)** *notum1*. **(G, H)** *wif1*. Digoxigenin whole mount in situ hybridization, glycol methacrylate emedded and sectioned 7 dpi kidneys. Scale bar in A = 10µm (applies to all panels).

**Supplemental figure 5.**
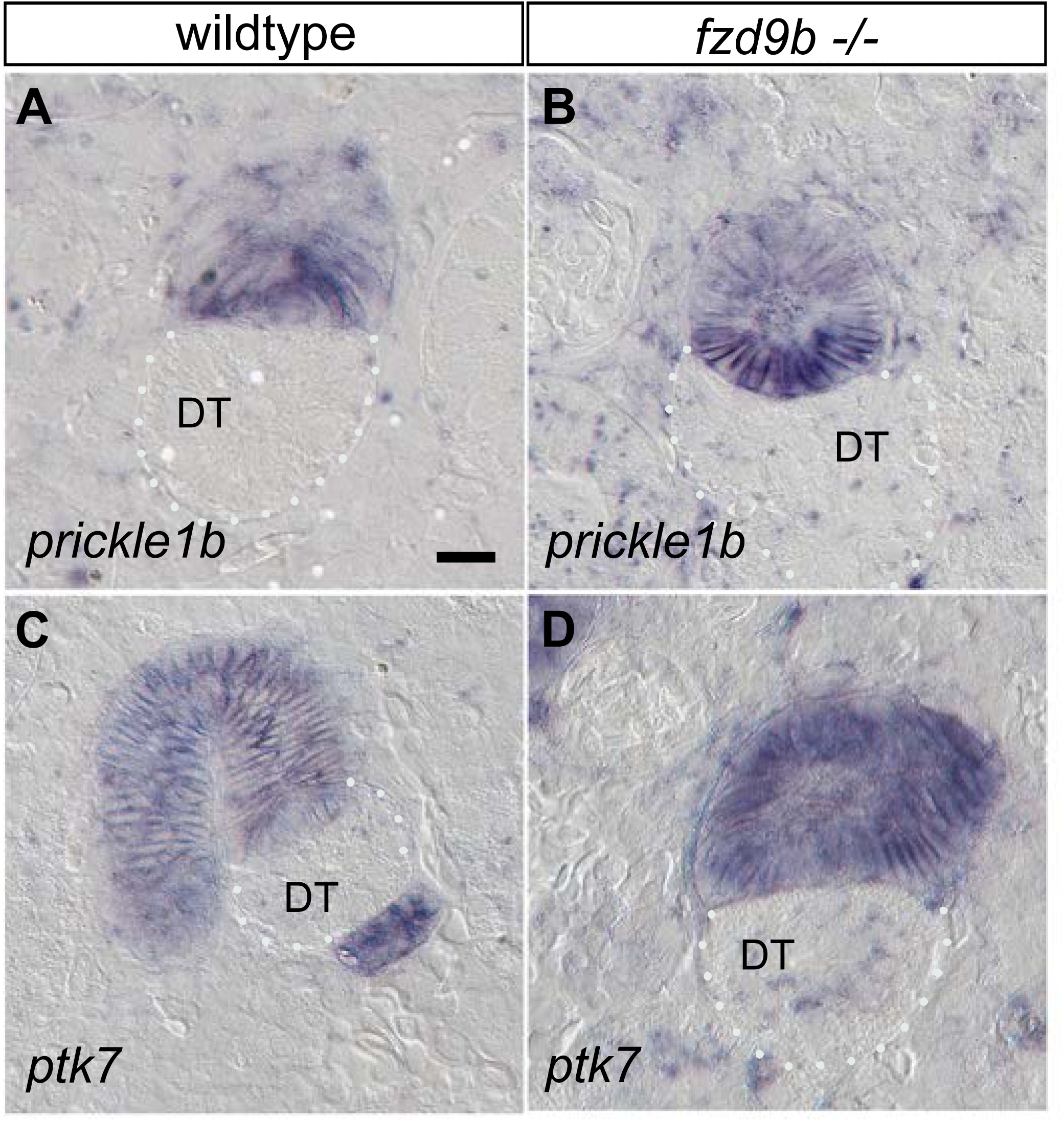
Expression of non-canonical Wnt pathway components in wildtype and *fzd9b* homozygous mutants. (A) *prickle1b* expression in wildtype 7 dpi new nephrons. (B) *prickle1b* expression in *fzd9b* -/- 7 dpi new nephrons. (C) *ptk7* expression in wildtype 7 dpi new nephrons. (D) *ptk7* expression in *fzd9b* -/- 7 dpi new nephrons. Digoxigenin whole mount in situ hybridization, glycol methacrylate emedded and sectioned 7 dpi kidneys. Scale bar = 10µm

**Supplemental figure 6.**
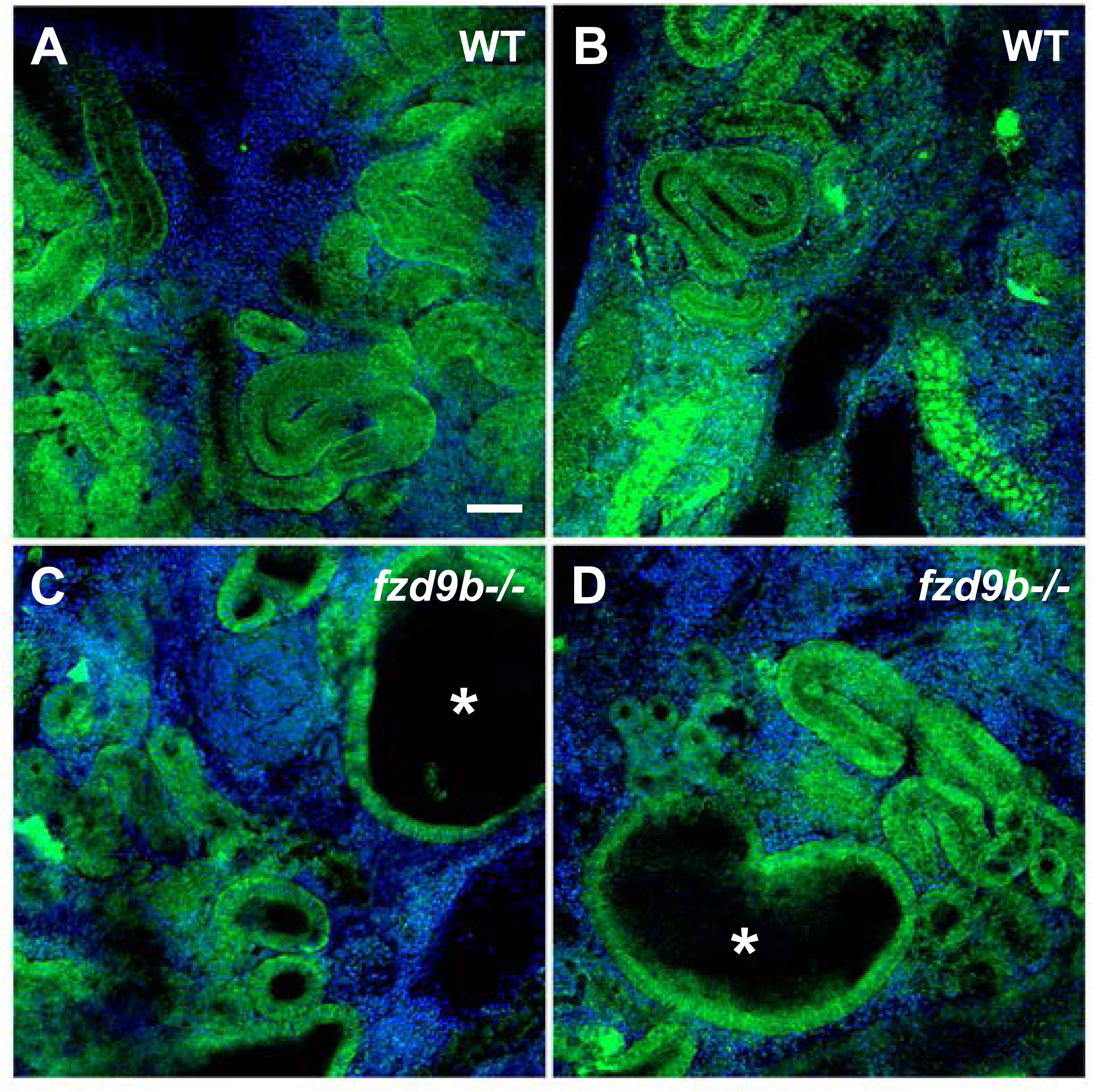
Two examples of tubule distension in *fzd9b* homozygous mutants at 14 dpi. (A,B) Wildtype (WT) confocal section of uninjured control kidney showing tubule sections in autofluorescence (green). **(C,D)** *fzd9b* -/- mutant kidney at 14 dpi shows distended tubules (*asterisk). Green = tubule cell autofluorescence, Blue = Hoechst DNA. Scale bar in A = 50 µm (applies to all panels).

## Supplemental movie legends

**Supplemental movie 1. Animated maximum intensity projection in support of** Figure 1A. Wildtype new nephron (green *Tg(lhx1a:eGFP)*) prior to fusion with the Distal Tubule stained with phalloidin (magenta) to highlight apical lumenal cell surfaces.

**Supplemental movie 2. Animated maximum intensity projection in support of** Figure 1B. Wildtype new nephron (green *Tg(lhx1a:eGFP)*) after fusion with the Distal Tubule stained with phalloidin (magenta) to highlight apical lumenal cell surfaces. Blue fluorescence is Hoechst-stained nuclei.

**Supplemental movie 3. Animated maximum intensity projection in support of** Figure 1G. Wildtype new nephron (green *Tg(lhx1a:eGFP)*) prior to fusion with the Distal Tubule showing invasive basal protrusions highlighted by eGFP expression. Proliferating cells are marked with EdU (magenta). Blue fluorescence is Hoechst-stained nuclei.

**Supplemental movie 4. Animated maximum intensity projection in support of** Figure 6D. *frizzled9b* homozygous mutant (*fzd9b* -/-) new nephron (green *Tg(lhx1a:eGFP)*) blocked from fusion with the Distal Tubule showing lateral misplacement of invasive basal protrusions highlighted by eGFP expression. Proliferating cells are marked with EdU (magenta). Blue fluorescence is Hoechst-stained nuclei.

**Supplemental movie 5. Animated maximum intensity projection in support of** Figure 7B. *frizzled9b* homozygous mutant (*fzd9b* -/-) new nephron (green *Tg(lhx1a:eGFP)*) blocked from fusion with the Distal Tubule stained with phalloidin (magenta) to highlight expanded apical cell surface area and failed apical constriction (compare with supplemental movie 1). Also note lateral growth of new nephron lumen parallel to the target DT. Blue fluorescence is Hoechst-stained nuclei.

**Supplemental movie 6. Animated maximum intensity projection in support of** Figure 7H. *frizzled9b* homozygous mutant (*fzd9b* -/-) new nephron (green *Tg(lhx1a:eGFP)*) blocked from fusion with the Distal Tubule stained with phalloidin (magenta) to highlight lateral lumen growth and displaced, constricted lumenal connection with the DT (compare with supplemental movie 2). Also note lateral growth of new nephron lumen parallel to the target DT. Blue fluorescence is Hoechst-stained nuclei.

